# Data Release: High-Resolution Imaging and Segmentation of P7 Mouse Tissue Microarchitecture Using FIB-SEM and Machine Learning

**DOI:** 10.1101/2024.09.05.611438

**Authors:** David Ackerman, Emma Avetissian, Christopher K. E. Bleck, John A. Bogovic, Michael Innerberger, Wyatt Korff, Wei-Ping Li, Zhiyuan Lu, Alyson Petruncio, Stephan Preibisch, Wei Qiu, Jeff Rhoades, Stephan Saalfeld, Malan Silva, Eric T. Trautman, Rebecca Vorimo, Aubrey Weigel, Zhiheng Yu, Yurii Zubov, CellMap Project Team

## Abstract

This report presents a comprehensive data release exploring the tissue microarchitecture of P7 aged mice using Focused Ion Beam Scanning Electron Microscopy (FIB-SEM) combined with machine learning-based segmentations of nuclei. The study includes high-resolution 3D volumes and nucleus segmentations for seven vital tissues—pancreas, liver, kidney, heart, thymus, hippocampus, and skin—from a single mouse. The detailed datasets are openly accessible onOpenOrganelle.org, providing a valuable resource for the scientific community to support further research and collaboration.

## Introduction

The postnatal day 7 (P7) developmental stage in mice represents a crucial period characterized by rapid tissue growth and complex cellular differentiation, setting the stage for organ development and functionality. To understand the microarchitecture of tissues at this critical juncture, advanced imaging techniques with nanometer-scale resolution and comprehensive cellular analysis are essential.

The CellMap Project Team employed Focused Ion Beam Scanning Electron Microscopy (FIB-SEM) to capture detailed 3D volumes of seven diverse tissues—pancreas, liver, kidney, heart, thymus, hippocampus, and skin—from a single P7 mouse. Utilizing machine learning-based segmentation, the team also generated nuclei segmentations within these tissues. This data release includes Transmission Electron Microscopy (TEM) micrographs and 3D X-ray volumes used for quality control screening and sample preparation.

These high-resolution FIB-SEM and X-ray volumes, along with their segmentations, are openly shared on OpenOrganelle.org, offering a unique opportunity to explore the intricate cellular landscapes and ultrastructural details. This report serves as an overview and an invitation for researchers across various fields to explore, analyze, and gain insights from these rich datasets, fostering collaboration and innovation within the scientific community.

### Sample Preparation

All experiments were approved by and performed in accordance with the guidelines of the Institutional Animal Care and Use Committee of Janelia Research Campus (protocol number 21-0206). Wild type mice (C57BL/6J strain, male from the Jackson Laboratory, ME) at postnatal day 7 (P7) were anesthetized with isoflurane and perfused with 0.13 M cacodylate buffer containing 2.5% paraformaldehyde and 2.5% glutaraldehyde (EMS, PA). The multiple tissues were dissected free and left to fix in the same fixative solution at 4°C for 24 hours. The samples were then sectioned into 300-µm-thick slices in 0.13 M cacodylate buffer using a Compresstome (Precisionary, MA). The slices were washed in cacodylate buffer (0.13 M), postfixed with 1% osmium tetroxide plus 0.8% potassium ferrocyanide in 0.13 M cacodylate buffer for 120 min at 0°C. After washing in distilled water, the slices were stained with 0.5% thiocarbohydrazide for 60 min and 2% osmium tetroxide for next 60 min at low temperature. After the heavy metal staining procedure, the samples were dehydrated and embedded in Durcupan resin (Sigma-Aldrich, MO) followed by polymerized at 60°C for 48 hours.

### Transmission Electron Microsocpy Imaging

The samples morphological quality was evaluated, Figure 1. After fixation, the samples were rinsed in 0.1 M cacodylate buffer and post-fixed in 1% osmium tetroxide for 1 hour at room temperature. The tissues were then sequentially dehydrated in a graded series of ethanol concentrations (30%, 50%, 70%, 90%, and 100%) followed by propylene oxide. Dehydrated samples were infiltrated with a mixture of propylene oxide and resin (1:1) for 1 hour, transferred to pure resin, and embedded in molds. The resin was polymerized at 60°C for 48 hours.

**Fig. 1.**
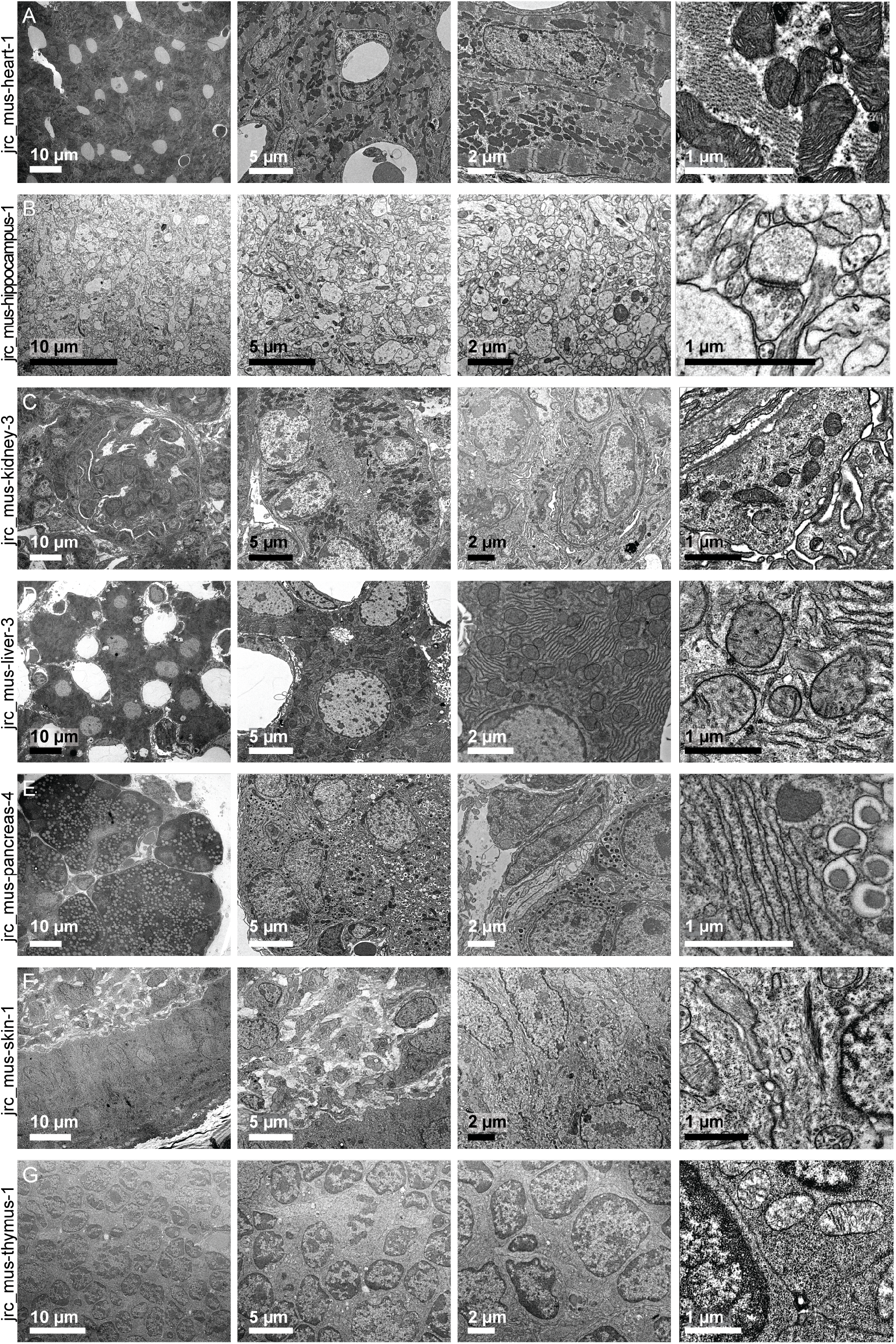
TEM quality checks for all seven diverse tissues—a) heart, b) hippocampus, c) kidney, d) liver, e) pancreas, f) skin, g) thymus—from a single P7 aged mouse.

Ultrathin sections (70–90 nm) were cut using a Leica UC7 ultramicrotome (Leica Microsystems) and collected on formvar/carbon coated 2 mm × 1 mm slot grids (Electron Microscopy Science Inc). No further staining was performed on these sections as one main goal of TEM imaging was to evaluate the quality of the en bloc staining for later FIB-SEM imaging. Grids with ultrathin sections were then loaded into an FEI Tecnai Spirit BioTWIN transmission electron microscope (TEM) equipped with a LaB6 filament (FEI Company). The TEM was calibrated and operated at 80 kV instead of the typical 120 kV for enhanced image contrast. Images were captured using a Gatan Oneview camera (Gatan Inc.) operated in mode 0. Images were first taken at a nominal magnification of 440 × from various locations on a sample to evaluate sample quality and to find areas for higher magnification imaging. Final images were taken at a nominal magnification of 2,900 × after focusing. The exposure time was set at 1 s, which results in no detectable sample drift. A 70 µm objective aperture was used for imaging. See Table 1 for hyperlinks and DOIs.

**Table 1.** Dataset and associated TEM DOIs.

| Dataset | Associated TEM |
| --- | --- |
| jrc_mus-heart-1 | 10.25378/janelia.26792734, 10.25378/janelia.26799169, 10.25378/janelia.26799283, 10.25378/janelia.26799292, 10.25378/janelia.26799298, 10.25378/janelia.26799304, 10.25378/janelia.26799370, 10.25378/janelia.26799379, 10.25378/janelia.26799388, 10.25378/janelia.26799403, 10.25378/janelia.26799412, 10.25378/janelia.26799418, 10.25378/janelia.26799424, 10.25378/janelia.26799427, 10.25378/janelia.26799433 |
| jrc_mus-hippocampus-1 | 10.25378/janelia.26799448, 10.25378/janelia.26799457, 10.25378/janelia.26799466, 10.25378/janelia.26799472, 10.25378/janelia.26799478, 10.25378/janelia.26799481, 10.25378/janelia.26799484, 10.25378/janelia.26799490, 10.25378/janelia.26799493, 10.25378/janelia.26799496, 10.25378/janelia.26799502, 10.25378/janelia.26799505, 10.25378/janelia.26799511, 10.25378/janelia.26799520, 10.25378/janelia.26799529, 10.25378/janelia.26799535, 10.25378/janelia.26799538 |
| jrc_mus-kidney-3 | 10.25378/janelia.26799568, 10.25378/janelia.26799577, 10.25378/janelia.26799583, 10.25378/janelia.26799586, 10.25378/janelia.26799592, 10.25378/janelia.26799595, 10.25378/janelia.26799598, 10.25378/janelia.26799604, 10.25378/janelia.26799607, 10.25378/janelia.26799610, 10.25378/janelia.26799772, 10.25378/janelia.26799775 |
| jrc_mus-liver-3 | 10.25378/janelia.26799781, 10.25378/janelia.26799784, 10.25378/janelia.26799790, 10.25378/janelia.26799799, 10.25378/janelia.26799802, 10.25378/janelia.26799811, 10.25378/janelia.26799823, 10.25378/janelia.26799832, 10.25378/janelia.26799838, 10.25378/janelia.26799844, 10.25378/janelia.26799853, 10.25378/janelia.26799856, 10.25378/janelia.26799886 |
| jrc_mus-pancreas-4 | 10.25378/janelia.26799925, 10.25378/janelia.26800003, 10.25378/janelia.26800033, 10.25378/janelia.26800060, 10.25378/janelia.26800078, 10.25378/janelia.26800102, 10.25378/janelia.26800111, 10.25378/janelia.26800135, 10.25378/janelia.26800144, 10.25378/janelia.26800252, 10.25378/janelia.26800291, 10.25378/janelia.26800300, 10.25378/janelia.26808271, 10.25378/janelia.26808421, 10.25378/janelia.26808424, 10.25378/janelia.26808439, 10.25378/janelia.26808448, 10.25378/janelia.26808451, 10.25378/janelia.26808460, 10.25378/janelia.26808472 |
| jrc_mus-skin-1 | 10.25378/janelia.26808502, 10.25378/janelia.26808511, 10.25378/janelia.26808514, 10.25378/janelia.26808517, 10.25378/janelia.26808541, 10.25378/janelia.26808544, 10.25378/janelia.26808547, 10.25378/janelia.26808553, 10.25378/janelia.26808556, 10.25378/janelia.26808565, 10.25378/janelia.26808577, 10.25378/janelia.26808583, 10.25378/janelia.26808586, 10.25378/janelia.26808589, 10.25378/janelia.26808595, 10.25378/janelia.26808622, 10.25378/janelia.26808637, 10.25378/janelia.26808640, 10.25378/janelia.26808667, 10.25378/janelia.26808670, 10.25378/janelia.26808694, 10.25378/janelia.26808703, 10.25378/janelia.26808715, 10.25378/janelia.26808721, 10.25378/janelia.26808730, 10.25378/janelia.26808754, 10.25378/janelia.26808757 |
| jrc_mus-thymus-1 | 10.25378/janelia.26808763, 10.25378/janelia.26808769, 10.25378/janelia.26808793, 10.25378/janelia.26808799, 10.25378/janelia.26808805, 10.25378/janelia.26808808, 10.25378/janelia.26808814, 10.25378/janelia.26808820, 10.25378/janelia.26808823, 10.25378/janelia.26808829, 10.25378/janelia.26808835 |

### Preparing Block for FIB-SEM Imaging

After re-embedding with Durcupan, a 3D micro-CT scan of the entire block was acquired using an XRadia Versa 510 Xray microscope (Zeiss Microscopy). The reconstructed tomogram from a micro-CT scan facilitated robust selection of Regions of Interest (ROIs) at micrometer or sub-micrometer resolution. Additionally, tomograms helped delineate the boundaries and orientation of tissue samples. The X-ray microscope was operated at 40 kV with a source power set at 3 W. Multiple rounds of micro-CT scans were performed on one resin block with different settings and experimental goals. Specifically, the 4 × objective lens was used to generate an overview tomogram. The 20 × objective lens was typically used to generate a tomogram for ROI picking. The number of projections for each micro-CT scan varies from 801 to 1,601.

Upon identifying a potential ROI, the sample block was remounted on a 1 mm copper post using Durcupan, ensuring contact with the metal-stained sample for improved charge dissipation, as previously described. The sample was oriented on the copper post to minimize the distance along the FIB milling direction. A vertical sample post, approximately 100 µm wide and 60–80 µm deep in the ion beam direction, was trimmed to the ROI using an ultramicrotome (EM UC7, Leica Microsystems). This trimming was performed iteratively, with X-ray tomograms collected after each step.

Once the desired ROI was trimmed, a thin layer of conductive material was applied. First, a 10 nm layer of gold was sputtered, followed by a 100 nm layer of carbon using a Gatan PECS 682 High-Resolution Ion Beam Coater. The coating parameters were 6 keV, 200 nA on both argon gas plasma sources, with a 10 rpm sample rotation and a 45° tilt.

### FIB-SEM Imaging

All seven samples were imaged at an 8 nm isotropic resolution using a customized Zeiss FIB-SEM system, which integrates a Zeiss Gemini SEM column with an FEI Capella FIB column (1,2). SEM imaging was performed with a 3 nA electron beam at a landing energy of 1.2 keV, operating at a raster frequency of 3.0 MHz, with an 8 nm x-y pixel size.

Both backscattered and secondary electron signals were captured via in-column detectors and optimally combined to enhance the signal-to-noise ratio. Sample surfaces were ablated layer by layer at an 8 nm Z-depth using a focused Ga^+^ beam at 30 keV and 13 nA beam current, operating in a closed-loop system. This FIB-ablation and SEM-imaging cycle was repeated over three to four weeks for each sample, with the acquisition time for these volumes ranging from three to four weeks.

### FIB-SEM Reconstruction

The sequence of acquired images formed a raw imaged volume, which was then processed for image registration and alignment. This process utilized Render, a volume assembly software pipeline designed to handle large-scale datasets (3). Render services from Trautman, Innerberger, Preibisch, and Saalfeld (2024), used in volume assembly for the adult Drosophila melanogaster ventral nerve cord (4), performed 2D stitching by extracting and matching similar features across overlapping images using Scale Invariant Feature Transform (SIFT) (5) following scan-correction for nonlinear speed-up of the detector. Subsequently, 3D alignment was achieved using a distributed implementation of a global optimization minimizing the distance between all corresponding SIFT features that were extracted across a range of 6 layers using a series of regularized affine models, producing isotropic 8 nm voxels. Finally, pixel intensities were adjusted using piece-wise affine filtering on a 8-by-8 sub-grid on the individual images, minimizing the intensity difference of pixels in overlapping image areas in 2D. Additionally, for the thymus dataset, Gauss-blurring with a radius of 50 px on 20 × downsampled images was used to estimate a background bias which was subsequently subtracted, and pixel intensities were also matched between adjacent z-layers. The final aligned stack consisted of an isotropic volume containing multiple complete cells, viewable in any arbitrary orientation, ensuring high-resolution 3D reconstructions and facilitating comprehensive analysis and visualization of the tissues’ microarchitecture. These volumes are available onOpenOrganelle.org, see Table 2 for hyperlinks and DOIs.

**Table 2.** Dataset, Registered X-ray, and FIB-SEM DOIs.

| Dataset | Registered X-ray | FIB-SEM |
| --- | --- | --- |
| jrc_mus-heart-1 | 10.25378/janelia.26530414, 10.25378/janelia.26813593, 10.25378/janelia.26813626, 10.25378/janelia.26813629 | 10.25378/janelia.24020919 |
| jrc_mus-hippocampus-1 | 10.25378/janelia.26530420, 10.25378/janelia.26813656, 10.25378/janelia.26813659, 10.25378/janelia.26813662 | 10.25378/janelia.24085176 |
| jrc_mus-kidney-3 | 10.25378/janelia.26530447, 10.25378/janelia.26813698, 10.25378/janelia.26813731 | 10.25378/janelia.24020799 |
| jrc_mus-liver-3 | 10.25378/janelia.26530450, 10.25378/janelia.26813767, 10.25378/janelia.26813770, 10.25378/janelia.26813818 | 10.25378/janelia.24051189 |
| jrc_mus-pancreas-4 | 10.25378/janelia.26530453, 10.25378/janelia.26813833, 10.25378/janelia.26820241, 10.25378/janelia.26820247 | 10.25378/janelia.23411843 |
| jrc_mus-skin-1 | 10.25378/janelia.26530459, 10.25378/janelia.26820253, 10.25378/janelia.26820256 | 10.25378/janelia.24085269 |
| jrc_mus-thymus-1 | 10.25378/janelia.26530462, 10.25378/janelia.26820259, 10.25378/janelia.26820265, 10.25378/janelia.26820271 | 10.25378/janelia.24085287 |

### FIB-SEM and X-Ray Registration

We took a sequential approach using Bigwarp (6) to manually register the X-ray tomograms to each other and the FIBSEM image data. First, the “Final Trim” was registered to the FIB-SEM using a thin-plate spline transformation. Next, the “Rough Trim” was registered to the “Final Trim” using an affine transformation. Finally, the “Overview” was registered to the “Rough Trim” also using an affine transformation, see Figure 2H. This implicitly brought all X-rays into alignment with the FIB-SEM image because concatenating transformations appropriately. For example, the “Overview” can be transformed into alignment with the FIB-SEM by concatenating the two affine transformations and the thinplate spline transformation. These volumes are available onOpenOrganelle.org, see Table 2 for hyperlinks and DOIs.

**Fig. 2.**
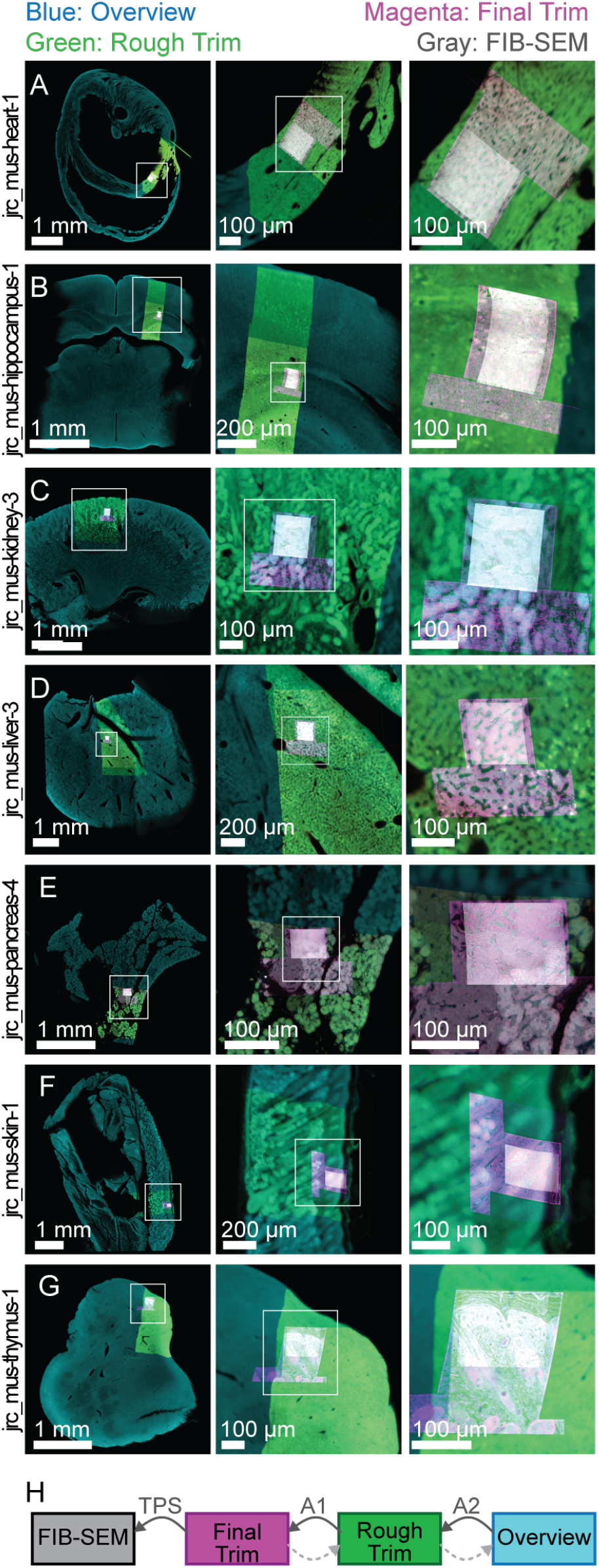
X-ray imaging of the large field of view and final trimmed block for each tissue sample—a) heart, b) hippocampus, c) kidney, d) liver, e) pancreas, f) skin,g) thymus—from a single P7 mouse. The white box indicates the region zoomed in on the middle and right panels. Included four overlaid images: the overview in blue, rough trim in green, final trim in magenta, and the FIB-SEM volume in gray. The left panel shows a large field of view overview of the prepared tissue, providing context for the selected region. The middle panel presents an intermediate zoom of the region of interest, and the right panel displays the final zoomed-in view of the selected region, highlighting the detailed preparation and orientation of the sample for subsequent FIB-SEM imaging. h) A graphical representation of the three image registrations performed to align the four images. A thin-plate spline (TPS) transformation aligns the Rough Trim to the FIB-SEM, and affines A1 and A2 align the Rough Trim to the Final Trim, and the overview to the Final Trim, respectively.

### Nuclei Segmentations

Nuclei segmentations were generated following a detailed protocol (7). The process began with preprocessing the images to enhance contrast and reduce noise. Next, the images were input into Cellpose (8–10), a deep learning-based segmentation algorithm, which was trained specifically for segmenting nuclei in electron microscopy datasets. Parameters for Cellpose were adjusted to optimize accuracy, including the diameter of the nuclei and flow threshold. Post-segmentation, the outputs were refined using manual correction tools to ensure precise segmentation. The trained models can be found on GitHub (janelia-cellmap/cellmap-models) (11), and the resulting nuclei segmentations are available on OpenOrganelle.org. See Table 3 for hyperlinks and DOIs. The results of this workflow are rendered in Figure 3.

**Table 3.** Dataset, Nucleus Segmentations, and Model DOIs.

| Dataset | Nucleus Segmentations | Model |
| --- | --- | --- |
| jrc_mus-heart-1 | 10.25378/janelia.25352203 | 10.25378/janelia.25352299 |
| jrc_mus-hippocampus-1 | 10.25378/janelia.25352212 | 10.25378/janelia.25352521 |
| jrc_mus-kidney-3 | 10.25378/janelia.25352215 | 10.25378/janelia.25352524 |
| jrc_mus-liver-3 | 10.25378/janelia.25352218 | 10.25378/janelia.25352545 |
| jrc_mus-pancreas-4 | 10.25378/janelia.25352224 | 10.25378/janelia.25352755 |
| jrc_mus-skin-1 | 10.25378/janelia.25352227 | 10.25378/janelia.25352773 |
| jrc_mus-thymus-1 | 10.25378/janelia.25352233 | 10.25378/janelia.25352779 |

**Fig. 3.**
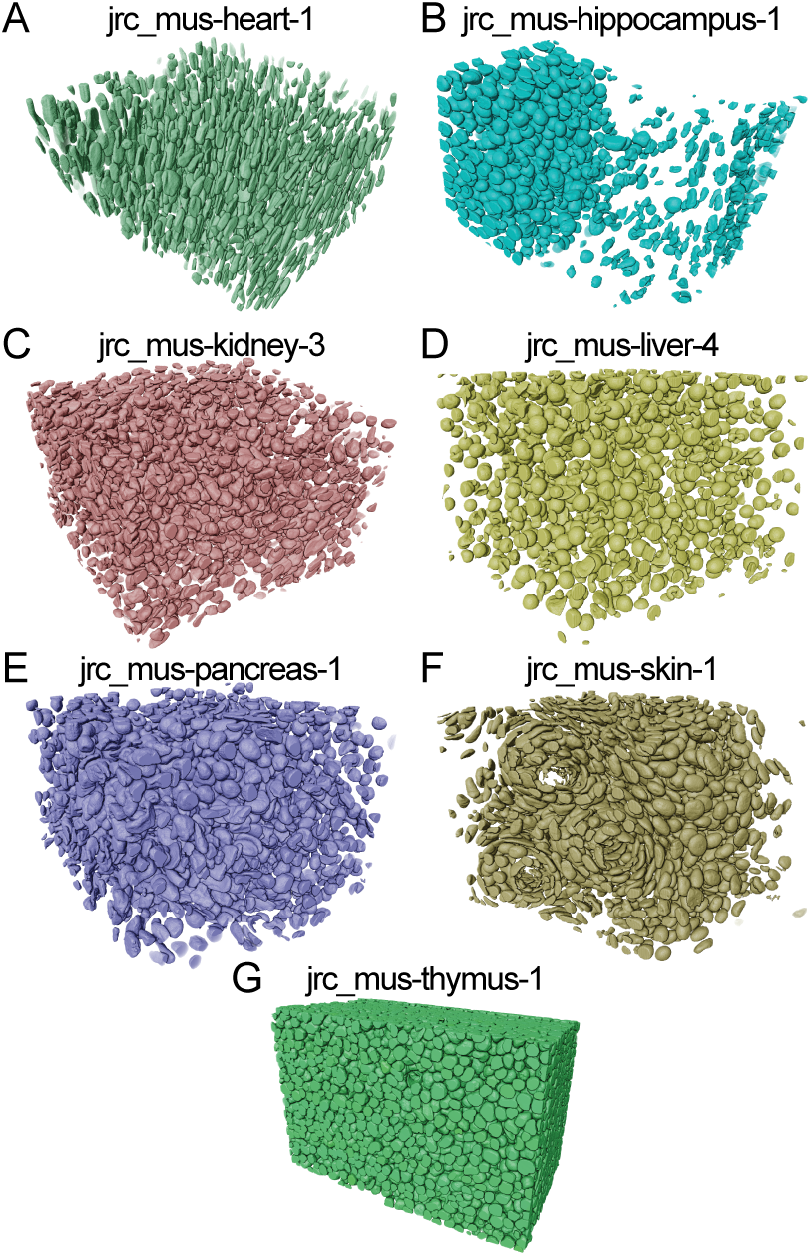
Nuclei segmentations of seven diverse tissues—a) heart, b) hippocampus,c) kidney, d), liver, e) pancreas, f) skin, g) thymus—from a single P7 aged mouse.

## Conclusions

The integration of FIB-SEM imaging with machine learning-driven segmentations has provided detailed datasets of the tissue microarchitecture of P7 aged mice. The dataset, openly available on OpenOrganelle.org and generated by the CellMap Project Team, serves as a valuable resource for researchers. This data release supports the scientific community by providing high-resolution 3D reconstructions and precise nuclei segmentations, facilitating detailed analysis and fostering further research and collaboration in developmental and computational biology.

## Data Availability

All of the data presented in this release can be found on OpenOrganelle.org. The trained Cellpose models are available on the CellMap GitHub: janelia-cellmap/cellmapmodels (11). Tables 1, 2, and 3 provides detailed information for each dataset.

## ACKNOWLEDGEMENTS

We would like to express our gratitude to the entire CellMap Project Team and CellMap Steering Committee for their dedication and expertise in generating and processing this comprehensive dataset. We also acknowledge AWS Open Data program for hosting the datasets and facilitating its accessibility to the broader scientific community. This work was supported by Howard Hughes Medical Institute, Janelia Research Campus.

## AUTHOR CONTRIBUTIONS

W.-P.L. carried out all sample collection. W.-P.L. and Z.L. prepared the samples for imaging and performed TEM imaging for quality check. X-ray microCT scans and trimming were done by Z.L. The collection of samples through block trimming was overseen by Z.Y. Regions of interest were chosen by C.K.E.B., W.-P.L., Z.L., and A.V.W. FIB-SEM images acquired by W.Q., C.K.E.B. oversaw. Post-processing was done by M.I., E.T.T., and S.P. The FIB-SEM and X-ray datasets were registered by J.B. A.V.W. and E.A. trained Cellpose models for nuclei segmentations. J.R. generated predictions using the trained models. A.V.W. and A.P. manually proofread and cleaned the predictions. D.A. generated meshes for the segmentations. R.V. and Y.Z. published the datasets to OpenOrganelle. A.V.W., J.B., W.-P.L., M.I., M.S., Z.Y., S.S., and S.P. wrote the manuscript. The rest of the CellMap Project Team contributed to general development and review of the project. W.K., A.V.W., and CellMap Project Team conceptualized the project. W.K. provided funding. A.V.W. managed the project.

The CellMap Project Team during this time consisted of: David Ackerman, Emma Avetissian, Davis Bennett, Marley Bryant, Hannah Nguyen, Grace Park, Alyson Petruncio, Alannah Post, Jacquelyn Price, Diana Ramirez, Jeff Rhoades, Rebecca Vorimo, Aubrey Weigel, Marwan Zouinkhi, Yurii Zubov. Misha Ahrens, Christopher Beck, Teng-Leong Chew, Daniel Feliciano, Jan Funke, Harald Hess, Wyatt Korff, Jennifer Lippincott-Schwartz, Zhe J. Liu, Kayvon Pedram, Stephan Preibisch, Stephan Saalfeld, Ronald Vale, and Aubrey Weigel were part of the CellMap Steering Committee.

